# Root growth promotion by *Penicillium melinii*: mechanistic insights and agricultural applications

**DOI:** 10.64898/2025.12.05.692050

**Authors:** Laura Gutierrez-Manso, Iván Devesa-Aranguren, Carlos M. Conesa, Gonzalo Monteoliva-García, Sandra González-Sáyer, Alberto Lozano-Enguita, Francesco Blasio, Lydia Ugena, José Nolasco, Ana Vázquez-Mora, Camila C.B. Levy, Enrique Ariel Otero, Patricia Fernández-Calvo, Miguel Ángel Moreno-Risueno, Ivan Petrik, Aleš Pěnčík, María Reguera, Sara González-Bodí, Jaime Huerta-Cepas, Soledad Sacristán, Juan Carlos del Pozo, Javier Cabrera

## Abstract

- This study characterizes *Penicillium melinii*, an endophytic fungus isolated from *Arabidopsis thaliana* roots, as a plant growth-promoting fungus with potential use as a model to study root development and as a biostimulant for sustainable agriculture. Although endophytes are known to promote plant growth, the underlying molecular mechanisms often remain poorly understood. Here, we aimed to elucidate how *P. melinii* enhances root system development and to assess its applicability across different crops.
- Phenotypic assays were conducted in Arabidopsis, quinoa and tomato under *in vitro*, greenhouse and field conditions. Root architecture and biomass were quantified using image-based phenotyping. Transcriptomic and phytohormone profiling assessed plant responses, and fungal genome sequencing coupled with secretome analysis was used to identify candidate effectors and metabolic traits.
- *P. melinii* consistently promoted root growth and increased plant biomass across species and environments, both *in vitro* and in the greenhouse. In tomato field trials, this translated into a significant increase in yield. The fungus colonized root surfaces without vascular penetration and triggered a mild transcriptomic response: early activation of stress-response genes followed by their attenuation and sustained upregulation of auxin-related pathways. Notably, the interaction modulates the *SLR-ARF-LBD* pathway and the number of pre-branch sites probably through increased auxin signalling in the oscillation zone. Additional hormonal changes were limited and mainly associated with the attenuation of the plant response to microorganisms.
- *P. melinii* enhances lateral root formation through a subtle molecular and metabolic dialogue with the host plant, underscoring its relevance as a model for studying root developmental plasticity. Its strong and reproducible growth-promoting effect, demonstrated with different fungal strains and under controlled and field conditions, supports its potential as a biostimulant for sustainable crop production.

## Introduction

Endophytic microorganisms, including fungi and bacteria, represent an important model for understanding plant–microbe interactions at the molecular levels. Their ability to influence root architecture and overall plant development provides a unique opportunity to explore the fundamental mechanisms underlying the plant-microorganism interaction and the root development (Gifford et al., 2024). Moreover, in the context of sustainable agriculture, these studies offer insights into how natural associations can be leveraged to reduce reliance on synthetic agrochemicals (Kumar et al., 2017; Baron & Rigobelo, 2021).

Recent advances highlight the need to unravel the molecular basis of plant–microbe interactions to optimise the use of endophytes as plant growth promoting (PGP) microorganisms. Despite progress, major gaps remain in understanding the complex signalling networks and metabolic exchanges between hosts and their microbial partners. The application of ‘omics’-based technology could help comprehend the complex associations existing between plants and their endophytes and can be a promising tool for sustainable environmental development (Anand et al., 2023). Addressing these gaps will enable innovative biotechnological strategies to advance sustainable agriculture and improve crop performance (Pereira et al., 2025; Kaur et al., 2023).

Endophytes influence plant development through multiple mechanisms, including modulation of phytohormone signalling, activation of stress-responsive pathways, and production of secondary metabolites (Fu et al., 2025; Vázquez-Chimalhua et al., 2021; Mora et al., 2023; Jensen et al., 2024; He et al., 2023). Auxin-related processes are particularly relevant for positive endophytes functions, as they can stimulate indole-3-acetic acid (IAA) synthesis and alter the expression of genes involved in auxin transport and signalling, such as *PINs*, *ARFs*, and *IAA14*, ultimately promoting lateral root (LR) formation and modifying root architecture (Fu et al., 2025; Sun et al., 2020). Some endophytes also trigger downstream cascades involving Lateral Organ Boundaries Domain (LBD) transcription factors and other regulators, occasionally bypassing canonical auxin receptors like TIR1/AFB2 (Jahn et al., 2021). In addition to hormonal modulation, volatile organic compounds (VOCs) such as DMHDA, acetoin, and sesquiterpenes influence root architecture through auxin, cytokinin, and ethylene signalling, activating hormone-responsive genes and those linked to cell wall remodelling and response to other organisms (Ditengou et al., 2015; Jiang et al., 2021). In addition to promote growth, endophytes further enhance resilience by upregulating stress tolerance genes and improving nutrient transport, and some activate alternative signalling routes, diversifying plant responses depending on the microorganism and environment (Gonin et al., 2023; Sun et al., 2020). Collectively, these findings highlight endophytes as dynamic regulators of plant growth and stress adaptation.

Among endophytic fungi, *Penicillium* species are increasingly recognised as important endophytes that colonise plant tissues without causing harm, establishing mutualistic interactions that enhance host fitness (Toghueo & Boyom, 2020). They promote growth through phytohormone production (e.g., gibberellins, IAA) (Leitão & Enguita, 2015) and nutrient solubilisation (Babu et al., 2015). *Penicillium* strains also mitigate abiotic stress by modulating hormonal balance and antioxidant systems (Leitão & Enguita, 2015). Strains such as *P. chrysogenum* and *P. simplicissimum* have been shown to improve crop productivity under stress conditions (Sun et al., 2025). Furthermore, these fungi produce diverse secondary metabolites with relevance in multiple sectors (i.e pharmaceutical or ecological), reinforcing their potential as bioinoculants in sustainable agriculture (Toghueo & Boyom, 2020). However, despite these benefits, the molecular mechanisms underlying *Penicillium*-mediated growth promotion remain poorly characterised. High-quality genomic resources for endophytic *Penicillium* species are scarce, limiting our ability to link functional traits to genetic determinants.

Here, *Penicillium melinii* is characterized as a novel endophytic fungus with strong root growth-promoting activity across model and crop species. *P. melinii* induces a subtle transcriptomic and hormonal response, suggesting a unique interaction strategy. The first mechanistic insights into this species were provided through transcriptomic, hormonal profiling, genome sequencing and *in silico* secretome analyses. The growth-promoting effect of *P. melinii* is conserved across multiple strains and validated under greenhouse and field conditions. Collectively, these findings establish *P. melinii* as a promising biostimulant and lay the foundation for its application in sustainable agriculture.

## Materials and Methods

### Plant Material, fungal strains and growth conditions

Fungal isolates screened *in vitro* were obtained from a collection of endophytes isolated from wild Arabidopsis population (García et al., 2013). Additionally, three strains of *Penicillium melinii* (CBS139142, CBS218.30, CBS285.65) were sourced from the Westerdijk Fungal Biodiversity Institute (https://wi.knaw.nl/).

A transgenic reporter line of *P. melinii* ‘isolate 2’ overexpressing GFP was generated for this work. Briefly, spores of *P. melinii* were obtained from 7-day-old Potato Dextrose Agar (PDA;Condalab) cultures, adjusted to OD₆₀₀ = 0.25–0.35 in induction medium containing acetosyringone and incubated for 2–6 h. After this time, fungal spores (10⁶ ml⁻¹) were mixed with *Agrobacterium tumefaciens* strain AGL1 carrying the plasmid pLM2 containing the fluorescent protein *eGFP* guided by the *β-tubulin* promoter (1:1) and co-cultivated on induction medium membranes for 48 h at 22 °C. Membranes were then transferred to PDA plates with hygromycin (100 µg ml⁻¹) and cefotaxime (200 µg ml⁻¹). Hygromycin-resistant colonies were subcultured on selective PDA plates after 7 days.

All fungal strains were maintained on PDA at room temperature and refreshed weekly.

*Arabidopsis thaliana* seeds used in this study included the Col-0 wild type, mutant lines *slr*, *arf7/arf19*, and *lbd16/lbd18*, as well as transgenic reporter lines *DR5:LUC;Col-0* (Moreno-Risueno et al., 2010), *DR5:LUC;arf7/arf19* (*DR5::LUC* was introgressed into mutant backgrounds arf7/arf19 by crossing), *proCYCB1:GFP* and *Lit6b-mCherry* (Elsayad et al., 2016). For greenhouse pot and rhizotron experiments and field trials, *Solanum lycopersicum* L. ‘Moneymaker’, ‘Optima’ and ‘N-283’ varieties were employed, respectively. *Chenopodium quinoa* willd cultivar Bastille was provided by the company Algosur S.L. (Lebrija, Spain) for the rhizotron experiments.

- *In vitro* assays: Arabidopsis plants were grown *in vitro* on square Petri dishes (12 × 12 cm) containing 1% agar (BD) and half-strength Murashige and Skoog (Caisson Labs; Murashige and Skoog, 1962) medium without vitamins or sucrose, supplemented with 300 µM KH₂PO₄ (Duchefa). Seeds were surface-sterilised in 30% bleach with 0.1% Triton X-100 (Sigma-Aldrich) for 15 min, rinsed five times with sterile distilled water, stratified for 2 days at 4 °C in darkness, and transferred to the D-root system (Silva-Navas et al., 2015) for 7 d. For Arabidopsis–fungus interaction tests, fungal isolates were inoculated onto identical plates with five 3-µL drops of spore suspension (10³ spores mL⁻¹) placed 3 cm from the seedling transfer area. After 4 days of fungal growth (7 days post-Arabidopsis sowing), five seedlings were transferred to each plate. Co-cultivation lasted 3-7 days before parameter quantification depending on the experiment.
- Greenhouse trials: For pot experiments in the greenhouse, tomato seeds were germinated in coarse vermiculite for 15 days and transplanted into 4 × 4 cm pots containing soil (amended with 0.073 g L⁻¹ single superphosphate and 3.018 g L⁻¹ of nitrogen and potassium). Three days after transplanting, 1 mL of spore suspension (10^7^ spores mL⁻¹) of the respective *P. melinii* strain, or water for the mock treatment, was applied adjacent to each stem. Plants were grown under controlled greenhouse conditions for 15 days before measuring the fresh weight of shoots and roots. For rhizotron experiments, tomato seeds were surface-sterilised as described above, germinated on moist sterile filter paper at 28 °C in darkness for 3–4 d, and seedlings with radicles ∼1 cm were transferred to rhizotron boxes filled with a 1:1 mixture of blond and black peat (Floratorf) at maximum water capacity. Three days after transplanting, 1 mL of spore suspension (10⁶ spores mL⁻¹) of *P. melinii* ‘Isolate 2’, or water for mock controls, was applied adjacent to each stem. Rhizotrons were covered with opaque black plastic, to keep the root system in darkness, and positioned vertically at a 50° angle. Root systems were imaged at multiple time points. Quinoa seeds were sterilised similarly and germinated directly in rhizotrons.
- Field trial: this assay was conducted by SynTech Research located in experimental fields in Las Cabezas de San Juan (Sevilla, Spain). A randomised complete block design was employed, comprising eight replications per treatment and an elemental plot size of 3.45 m² (10 plants per plot). Each plant received either tap water or a spore suspension (2 × 10⁷ spores mL⁻¹). Treatments were applied manually, directly to the soil at the base of the plant stem, 10 days after transplanting, using an application volume of 200 mL per plant. The fresh and dry weight of the aerial biomass were recorded 21 days after treatment. Three months after the inoculation, tomatoes were harvested and the yield was calculated as kilograms per hectare (kg ha⁻¹).

### Phenotyping and Imaging

- Quantification of root system parameters: For Arabidopsis *in vitro* assays, primary root length and LR number were measured from the transfer point using Fiji software (Schindelin et al., 2012). In rhizotron experiments, images were acquired using the LemnaTec phenotyping software^©^. Root segmentation employed a convolutional neural network trained on 15 images under seven exposure levels, iteratively refined over eight rounds, achieving ∼99% accuracy (loss = 0.05). Post-segmentation, background pixels were removed via erosion and dilation steps using LemnaTec software.
- Auxin signalling analysis with the *DR5:LUC* Reporter transgenic line: Auxin signalling was assessed in *DR5:LUC* lines (either in Col-0 or *arf7/arf19 genetic* backgrounds) following co-cultivation with fungal isolates. After 5 days of interaction, seedlings were imaged using a NightOwl II (Berthold Technologies) system equipped with a CCD camera after applying 1 mL of 2.5 mM luciferin. Luminescence in the oscillation zone (OZ) and pre-branch sites (PBS) was captured with a 300 s exposure. Maximum luminescence intensity in OZ was quantified for *DR5:LUC*;*arf7/arf19*, and PBS counts were recorded for *DR5:LUC*;Col-0, in both cases using IndiGO™ software (Berthold Technologies).
- Confocal laser scanning microscopy: Meristem activity was visualised using a Leica SP8 microscope. GFP-tagged cells in *proCYCB1:GFP* roots and GFP-expressing fungal lines were detected at 488/510 nm (excitation/emission). Cell walls were stained with 5 µg mL⁻¹ propidium iodide (PI) and detected at 535/617 nm.

### Statistical analysis

Differences between inoculated and mock plants were tested using Student’s t-test (P < 0.05). Grubbs’ test was used to detect outliers. The effect size representing the difference between the groups was computed as Cohen’s *d*, obtained by dividing the mean difference between groups by the pooled standard deviation. Root growth in rhizotrons was analysed with a linear mixed-effects model (Sample as random effect growth and time fixed as effect). Marginal and conditional R² values were calculated. Analyses were performed in R using the lme4 package (https://cran.r-project.org/web/packages/lme4/index.html). For hormone profiling analysis statistical significance was assessed using the Wilcoxon-Mann-Whitney U test. P-values were transformed to a -log_10_ scale for visualization in volcano plots. A significance threshold of α = 0.05 was applied, and metabolites exceeding this threshold are indicated above the horizontal dashed line.

### Biochemical and transcriptomic analyses

- Quantification of indole derivatives: Indole compounds were quantified using the Salkowski colorimetric method (Gang et al., 2019) by measuring absorbance at 530 nm. *P. melinii* was cultured in modified Czapek–Dox medium supplemented with varying L-tryptophan concentrations at 28 °C, 200 rpm, in darkness for 7 d. Indole compound levels were determined by interpolation against an IAA standard curve.
- RNA extraction, sequencing and analysis: Total RNA was extracted from roots of ‘isolate 2’-inoculated and mock plants at 3 and 6 days post inoculation (dpi). Both aerial and root tissues were collected, frozen in liquid nitrogen, and ground into powder. Approximately 50-100 mg from three biological replicates was processed using the RNeasy Plant Mini Kit (QIAGEN). RNA quality was assessed with NanoDrop. Twenty-four samples were sequenced by BGI’s Illumina platform (China). Differentially expressed genes (DEGs) were identified using DESeq2 v1.40.1 (Padj < 0.05; log₂FC > 0.5 or < −0.5; Love et al., 2014). Venn diagrams were generated with Venny software (Oliveros, 2007), and gene term enrichment analysis was performed using Metascape (Zhou et al., 2019).
- Phytohormone profiling: Root and shoot tissues from mock and *P. melinii* ‘isolate 2’-inoculated plants were collected at 3, 6, and 12 dpi, frozen, and lyophilised. Five biological replicates per treatment were analysed for phytohormone-related metabolites following the protocol described in Simura et al. (2018). Briefly, ∼2 mg DW was extracted in 1 mL of aqueous 60% (v/v) acetonitrile. A mixture of stabile isotopic labelled internal standards was added to ensure the accurate plant hormone quantification. The crude extract was centrifuged. The supernatant was purified with Oasis ® HLB 1 mg/1 cc solid phase extraction cartridge (Waters, Milford, CT USA). The flowthrough fraction was collected and evaporated under a reduced pressure. The samples were reconstituted in aqueous 25% (v/v) acetonitrile and 1/8 portion of the sample was injected into Acquity UPLC® CSH^TM^ C18 150 × 2.1 mm, 1.7 µm chromatographic column (Waters, Milford, CT USA). The chromatographic separation was performed using Acquity I-Class system (Waters, Milford, CT USA). The separated sample was analysed using a triple quadrupole Xevo TQ-XS system (Waters, Manchester, UK) equipped with electrospray ionization. Data were processed with MassLynx v4.2 software (Waters, Manchester, UK). The plant hormones were quantified using the isotope dilution method.

### Genomic analyses

- Whole-Genome sequencing and assembly: Genomic DNA from *P. melinii* ‘isolate 2’ was extracted using the Qiagen DNeasy kit. Sequencing was performed by Macrogen Inc. on the Illumina NovaSeq® 6000 platform. Reads were quality-checked using FastQC v.0.12.0 (Andrews, 2010) and MultiQC v.1.21.0 (Ewels et al., 2016), trimmed using Trimmomatic v.0.38 (Bolger et al., 2014), and re-checked with FastQC/MultiQC to confirm post-trimming quality and re-evaluated for quality. Assemblies were generated using SPAdes v3.15.2 (careful mode, k-mers: 21–127) (Prjibelski et al., 2020) and MaSuRCA v4.1.1 (default) (Zimin et al., 2013). Genome assembly contiguity and general quality metrics were evaluated using QUAST v.5.2.0 (Gurevich et al., 2013) and completeness was assessed with BUSCO v.5.7.1 (Manni et al., 2021).
- Functional Annotation: Initially, repetitive sequences were masked with RepeatModeler v.2.0.1 (Smit et al., 2018) and RepeatMasker v.4.1.2-p1 (Smit et al., 2015). GeneMark-ES v.4.30 (Lomsadze et al., 2005) was then used for gene prediction. Functional annotation of predicted genes was carried out using the following tools: BlastKOALA (Kanehisa & Sato, 2020) for KEGG orthology assignments, EggNOG-mapper v.2.1.12 (Huertas-Cepas et al., 2016) for COGs (Clusters of Orthologous Genes; http://eggnog-mapper.embl.de/), and InterProScan v.5.66-98.0 (Jones et al., 2014) for GO terms and protein domain architecture. Furthermore, the identification of secreted proteins was conducted using SignalP (Teufel et al., 2022) and TMHMM v.2.0c (Krogh et al., 2001) tools, while the removal of ER-retained proteins was achieved through the implementation of the ProSite PS00014 signature (Blum et al., 2021). Carbohydrate-active enzymes (CAZymes) were identified using CUPP v.2.1.0 (Barret et al., 2020) and dbCAN3 v.12.0 (Zheng et al., 2023). Proteases were detected via BLASTP (Camacho et al., 2009) against the MEROPS database (Rawlings et al., 2009), effectors with EffectorP v.3.0 (Sperschneider & Dodds, 2022), and transporters via BLASTP against TCDB (Saier et al., 2016). Finally, secondary metabolite clusters were predicted using antiSMASH v7.0 (Blin et al., 2021).
- *P. melinii* taxonomic identification: *P. melinii* ‘isolate 2’ taxonomic identification was performed following the methodology described by Houbraken et al. (2020). Partial sequences of *β-tubulin* (*BenA*), *calmodulin* (*CaM*), and the second largest subunit of *RNA polymerase II* (*RPB2*) were obtained using specific primers. These sequences were concatenated and aligned with the corresponding concatenated sequences from 94 species of *Penicillium* and related genera, including *Hamigera avellanea* as the outgroup. Sequences for each locus were downloaded from NCBI, and multiple sequence alignments were generated using MAFFT v7.505 (Katoh & Standley, 2013) with the ‘--auto’ parameter. These alignments were then trimmed using a custom python3 script and preliminary gene-specific phylogenetic trees were inferred using FastTree to assess the topology of each individual marker. For the concatenated dataset, alignments of the three loci were combined using the catfasta2phyml perl script (Nylander, 2010). Maximum Likelihood (ML) phylogenetic tree was then generated with the concatenated dataset using IQ-TREE v2.2.5 (Minh et al., 2020) and partitioned models for each marker. ModelFinder (Kalyaanamoorthy et al. 2017) was used to automatically determine the best-fitting substitution model for each partition, resulting in TNe+I+G4 for BenA and RPB2, and TIM3e+I+G4 for CaM gene. Branch support was assessed using 1,000 ultrafast bootstrap replicates (-bb 1000). The resulting concatenated tree was visualised and annotated using ETE v.4 (Huerta-Cepas et al., 2016), with leaf nodes classified according to series and section.
- Phylogenomic analysis: Outgroup for phylogenomic placement of *P. melinii* ‘isolate 2’ was conducted using 607 publicly available genomes of the genus *Penicillium* from the NCBI and *Trichoderma reesei* as outgroup. For phylogenomic tree reconstruction, we followed the approach described by Badet & Croll (2025). Briefly, single copy orthologs were identified with BUSCO v.6.0.0 (Simão et al., 2015) using Ascomycota database (odb10). Of the 1706 BUSCO genes analysed, those with ≥50% occupancy across Penicillium genomes were retained. From these, 100 single-copy BUSCO genes amino acid sequences were randomly selected, concatenated using AMAS v1.0 (Borowiec, 2016), and aligned with MAFFT v7.526 using the parameters --genafpair --maxiterate 1000. Poorly aligned regions were trimmed with TrimAl v1.5 (Capella-Gutiérrez et al., 2009) using -automated1 option. Maximum likelihood inference was conducted with IQ-TREE v2.2.0 (Minh et al., 2020) applying ModelFinder Plus (MFP) for model selection and MERGE for partition merging. Branch support was assessed using 1000 ultrafast bootstrap replicates (-bb 1000) and 1000 SH-aLRT tests (-alrt 1000). The *Penicilium* species tree was inferred with ASTRAL v5.7.8 (Zhang et al., 2018) and visualized using R packages phylotools (Zhang, J. et al., 2010), treeio (Wang et al., 2020), ggtreeExtra (Xu et al., 2021).

## Results

### The endophytic fungal ‘isolate 2’ from Arabidopsis enhances plant root system development

To identify a novel fungus that enhance root system development, a screening of a collection of endophytic fungi isolated from *Arabidopsis thaliana* (García et al., 2013) was performed. The selection was refined to include isolates capable of withstanding long-term freezing, exhibiting high sporulation rates, and growing on MS medium without sucrose. Eleven isolates were subsequently tested for their ability to promote root growth in Arabidopsis through *in vitro* interaction assays and primary root length and LR number were assessed as indicators of root system development (Figure S1). Up to four isolates exhibited a negative impact on both primary root elongation and LR formation (Figure S1A–B). In contrast, ‘isolate 2’ stood out, significantly increasing both the length of the primary root and the number of LRs (Figure S1A-B), resulting in a marked enhancement of the overall Arabidopsis root system (Figure 1A).

**Figure 1.**
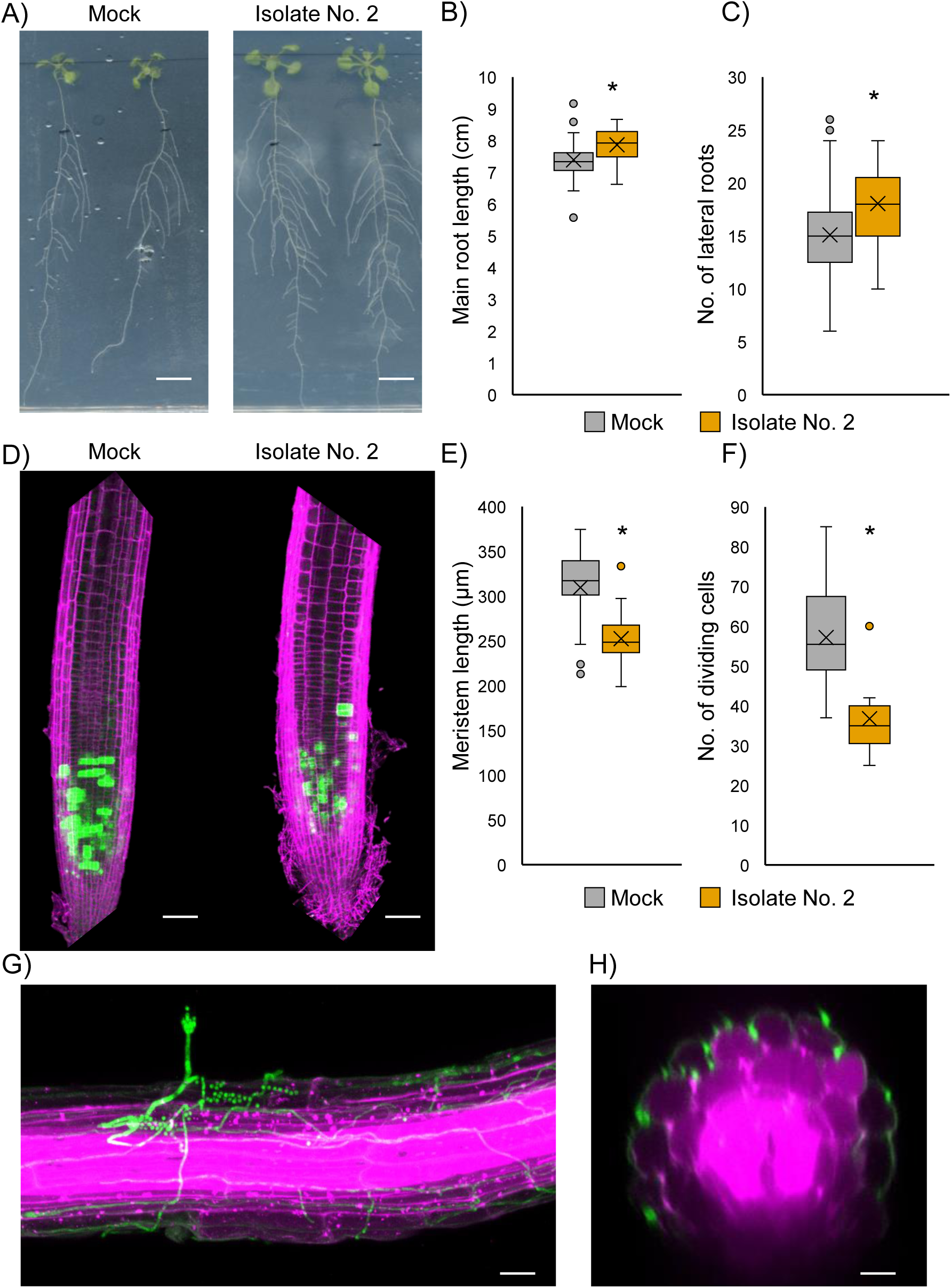
Fungal strain ‘isolate 2’ enhances root system development in Arabidopsis. (A) Representative images of Arabidopsis seedlings after seven days of incubation with fungal ‘isolate 2’. (B) Quantification of primary root length and (C) number of lateral roots, measured seven days after incubation. Scale bar: 1 cm. Measurements were taken from the root section that developed on the fungal-inoculated plate after seedling transfer. Asterisks indicate statistical significance (Student’s t-test, n > 28, P < 0.05). (D) Visualization of dividing cells in the root apical meristem in the *proCYCB1:GFP* Arabidopsis line, either inoculated or not inoculated with ‘isolate 2’. GFP (green) marks dividing cells; propidium iodide (pink) stains cell walls. (E) Quantification of root apical meristem length and (F) number of dividing cells in the *proCYCB1:GFP* line, measured three days after interaction with ‘isolate 2’ (Student’s t-test, n > 9, P < 0.05). (G–H) Representative longitudinal and transverse images showing GFP-expressing ‘isolate 2’ (green) colonizing the *Lti6b-mCherry* membrane marker Arabidopsis line (pink).

Previous research has described that artificial illumination on roots can *bias* the plant’s response to environmental stresses and interactions (Silva-Navas et al., 2015; Cabrera et al., 2022). Hence, the beneficial effect of this interaction was validated using the D-Root system (Figures B-C; Silva-Navas et al., 2015). Plant-to-plant variability was reduced under D-Root conditions, and the effect sizes for primary root length and LR number (Cohen’s d = 0.73 and 0.72, respectively) indicated a moderate to large impact of ‘isolate 2’ on these traits. These findings demonstrate that ‘isolate 2’ is capable of promoting root growth in Arabidopsis under *in vitro* conditions. Root meristem activity under inoculation with ‘isolate 2’ was assessed using the *proCYCB1:GFP* line, a cell division-reporter (Figures 1D–F). Root meristems from plants treated with ‘isolate 2’ were shorter (Figure 1E) and contained fewer dividing cells compared to those of non-inoculated controls (Figure 1F). These results suggest that root growth is driven by accelerated cellular differentiation and enlargement, rather than by an increase in meristem size, at least at these early stages of the interaction.

To verify the type of interaction between ‘isolate 2’ and the root, a transgenic fungal line expressing GFP under the β-tubulin promoter was generated (Isolate 2-GFP). Robust colonisation by ‘isolate 2-GFP’ was observed along the entire root length within the outer cell layers (Figure 1G), with hyphae penetrating between the epidermal cells (Figure 1H). No differences in colonisation were found between different root regions, and colonisation of the vascular cylinder was not detected.

### The root growth–promoting activity of ‘isolate 2’ extends to crops

To explore the conservation of the beneficial capacity of the fungus in an alternative interaction system, validation experiments under greenhouse conditions using different plant species, as quinoa and tomato, were performed. To monitor root system expansion more precisely over time, a rhizotron system was employed (Figure 2A). Quinoa seedlings were inoculated with ‘isolate 2’ (Figures 2A-C). A 15% faster growth increase in root area was observed 17 days post-inoculation compared to controls (Figures 2A-B), with a significant growth trend differentiating the treatments over time. The model demonstrated adequate explanatory capability, evidenced by a marginal R^2^ of 0.73 for the fixed effects and a conditional R^2^ of 0.80 when random effects were incorporated. With respect to the baseline timepoint, a significant increase in response to the inoculation was observed at 10 to 17 dpi, with the most pronounced effect occurring on the latter timepoint. The interaction terms indicated that the roots inoculated with the ‘isolate 2’ exhibited a growth trajectory analogous to that of the reference group at early dates but a statistically significant increase was observed 10 and 14 dpi, which was consistent with a group-specific enhancement at the final sampling timepoint (Figure 2B). This enhanced root development correlated with an increase in aerial biomass, assessed as leaf surface area from top and lateral views (Figure 2C). Again, the model demonstrated adequate explanatory power, with the fixed effects accounting for 71% of the variance (marginal R^2^=0.71) and the full model, incorporating random effects, explaining 81% (conditional R^2^=0.81). The interaction terms demonstrated that the Mock treatment exhibited a comparable temporal pattern at early dates, as observed in the preceding analysis. Nevertheless, plant inoculation with isolates demonstrated significantly elevated responses in comparison to the Mock treatment at 14 and 17 dpi, signifying a treatment-specific positive response at the latest sampling time (Figure 2C).

**Figure 2.**
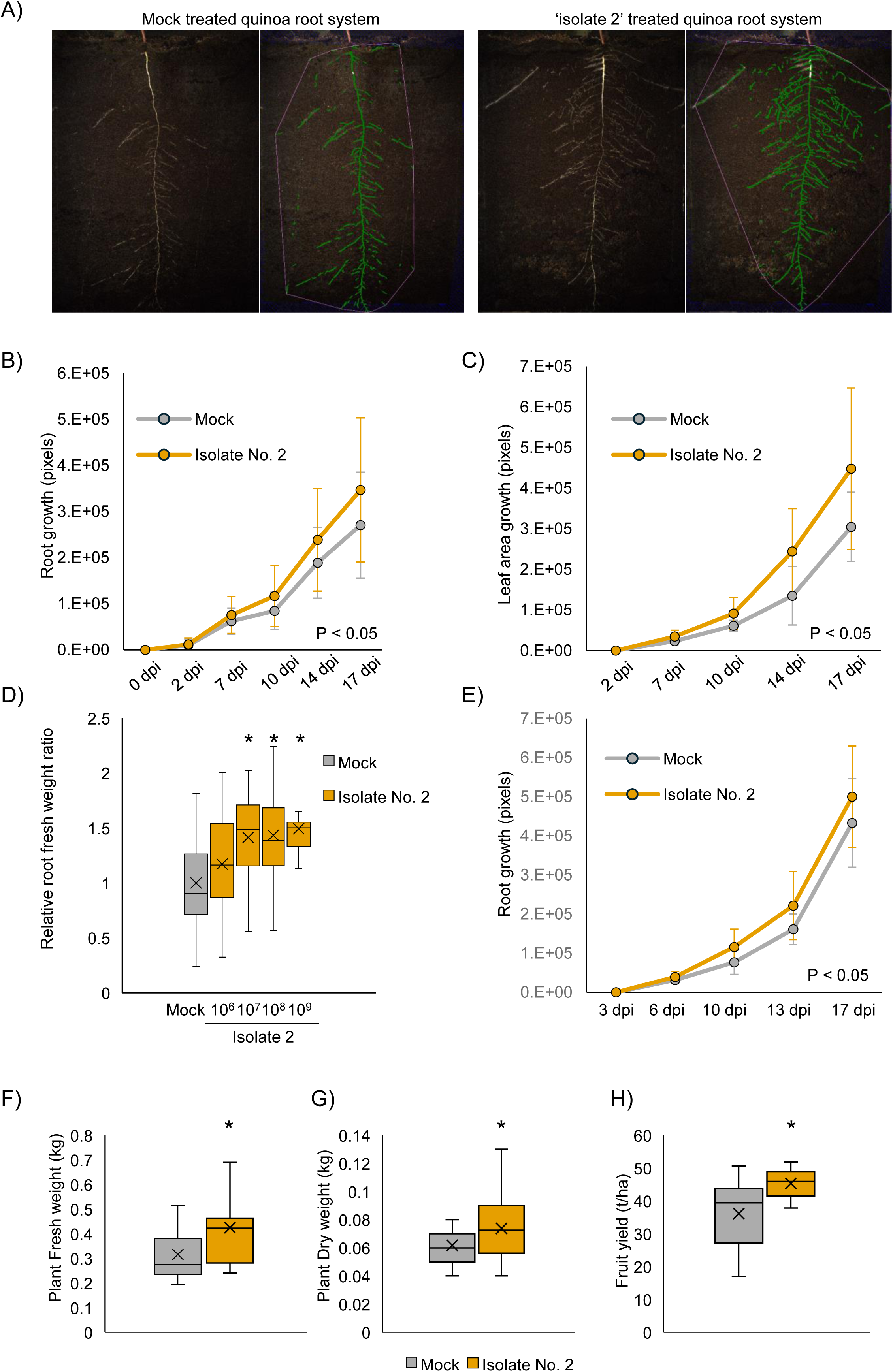
‘Isolate 2’ preserves its root growth-promoting activity in agricultural crops. (A) Representative images of quinoa root systems grown in rhizotrons with or without ‘isolate 2’. (B) Quantification of total root growth and (C) total leaf area in quinoa plants grown in rhizotrons and inoculated with ‘isolate 2’, measured over time (Generalized Linear Mixed Model, n = 15, P < 0.05). (D) Fresh weight of tomato plants grown in soil pots and inoculated with varying concentrations of ‘isolate 2’ spores. Asterisks denote statistically significant differences (Student’s t-test, n > 28, P < 0.05). (E) Quantification of total root growth over time in tomato plants grown in rhizotrons following inoculation with ‘isolate 2’ (Generalized Linear Mixed Model, n = 10, P < 0.05). (F) Fresh and (G) dry aerial biomass and (H) tomato fruit yield from field trials comparing plants inoculated or not with ‘isolate 2’ (Student’s t-test, n = 80, P < 0.05).

To extend this investigation to other agriculturally relevant species, twenty-day-old tomato seedlings were inoculated with varying doses of fungal spores (Figure 2D). Ten days post-inoculation, a dose-dependent increase in root fresh weight was observed, with the highest dose (10⁹ spores) resulting in a root system approximately 50% larger than that of control plants. The effect size at this maximum dose, calculated using Cohen’s d, was 1.25, indicating a strong positive impact of ‘isolate 2’ on root development. In rhizotrons, quantitative and qualitative analysis demonstrated a significant increase in root growth in tomato plants inoculated with ‘isolate 2’ compared to mock-treated controls (Figure 2E). Seventeen days after inoculation, inoculated plants exhibited on average 15% faster growth than non-inoculated plants, with a Cohen’s d of approximately 0.82 between 10- and 17-days post-inoculation, indicating a robust and meaningful effect (Figure 2E). In fact, the linear mixed model utilised for the analysis of root growth exhibited a high degree of explanatory power, with a marginal R² of 0.87 for the fixed effects and a conditional R² of 0.92 for the full model that incorporated random effects. Across the observed timepoints, there was a marked increase in response to inoculation relative to the baseline date, with the most significant increase observed at 17 dpi (Figure 2E).

In addition, the capacity of ‘isolate 2’ to enhance plant growth under natural conditions was assessed through field trials with tomato plants inoculated with this fungal strain. These trials resulted in a significant increase in the leaf biomass, both in fresh or dry weight, and in the fruit yield after harvesting (Figures 2F-H).

These findings confirm that ‘isolate 2’ not only stimulates root growth *in vitro* in Arabidopsis but also promotes root development in soil-grown tomato and quinoa plants under greenhouse conditions and of tomatoes growing in open field. This reinforces its potential as a promising biostimulant across species for agricultural applications.

### Phylogenetic analysis confirms ‘isolate 2’ as *Penicillium melinii*, a species with conserved growth-promoting traits

The identification and taxonomic placement of ‘isolate 2’ within the *Penicillium* genus were assessed using a phylogenetic approach (Figure 3D). The locus-based phylogenetic analysis strongly supported the placement of ‘isolate 2’ within a clade that exclusively includes *P. melinii*, with a bootstrap value of 100%. The *P. melinii* strains showing the highest sequence similarity were CBS 340.61, CBS 280.58, and CBS 285.65, all grouped in the same subclade belonging to the Exilicaulis section (Houbraken & Samson, 2011). These results confirmed that ‘isolate 2’ corresponds to the species *P. melinii* (Figure 3D). Additionally, a genetic marker similarity analysis based on the ITS region was performed by comparing nucleotide sequences with those retrieved from the NCBI GenBank database. The highest similarity (100% nucleotide identity) was observed with the sequence under accession number NR_077155.1, *P. melinii* FRR 2041 ITS region (TYPE material, as of 22/03/2024).

**Figure 3.**
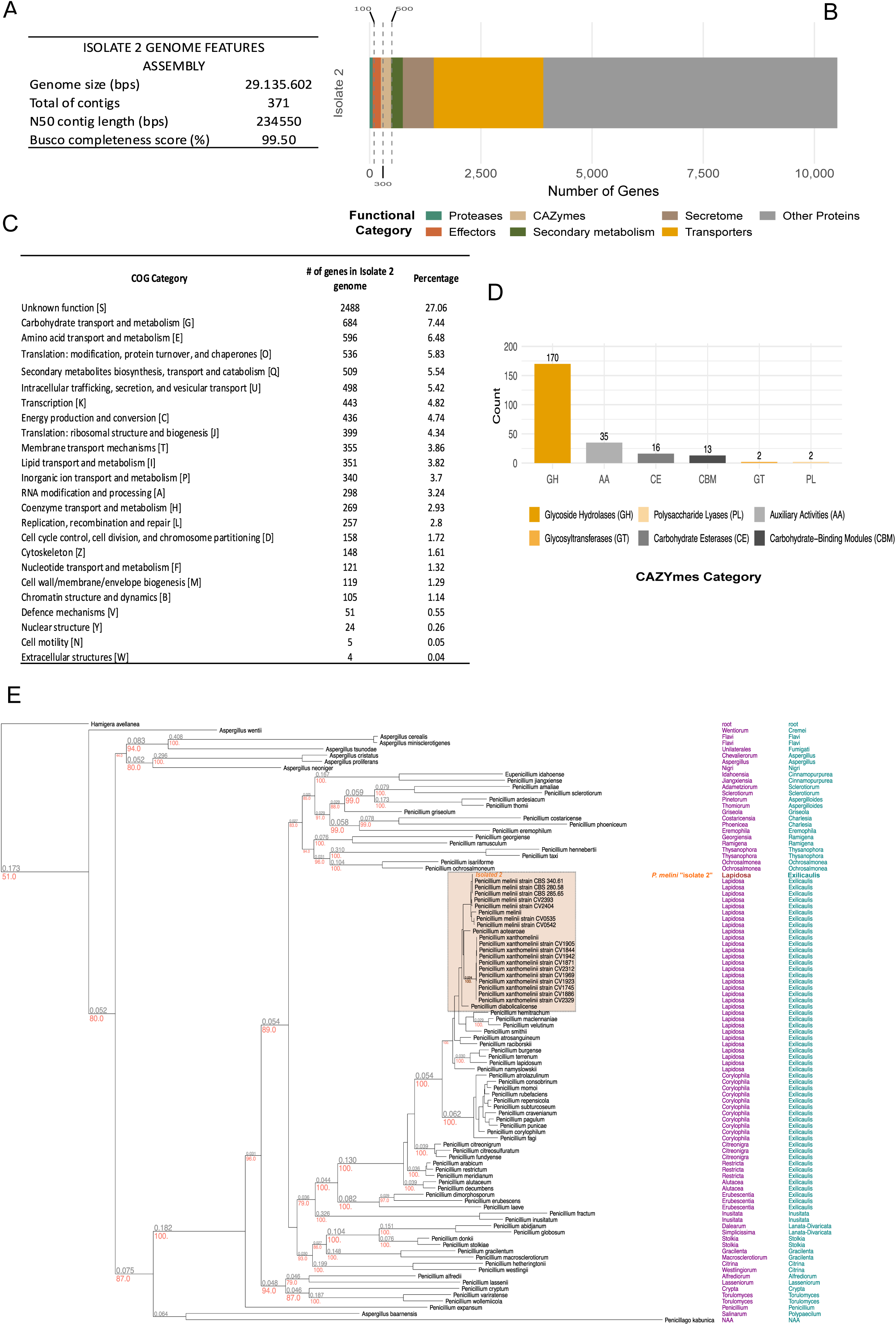
Genomic characterization and functional features of ‘isolate 2’. A) ‘isolate 2’ genome assembly statistics. (B) Functional groups associated with plant growth promotion, including CAZymes, proteases, and predicted effectors within the secretome. (C) Gene prediction and annotation results, showing coding sequence counts and functional categorization. (D) Number of CAZymes with different enzymatic activities found in the genome (E) Taxonomic placement of ‘isolate 2’ as *Penicillium melinii* based on phylogenetic analyses.

To determine whether the species *P. melinii* exhibits growth-promoting properties beyond those of ‘isolate 2’, three additional strains from international public repositories were evaluated: CBS139142 (South Africa), CBS218.30 (USA), and CBS285.65 (England) (Figure 5A). A comparable increase in root and shoot fresh weight of tomato plants was observed 10 days post-inoculation with either ‘isolate 2’ or these strains (Figures 5B-C). This result provides compelling evidence that the growth-promoting capacity of *P. melinii* is a conserved, species-level trait rather than an isolate-specific phenomenon. Such consistency across genetically and geographically distinct strains substantially reinforces its potential as a broad-spectrum biostimulant for sustainable crop production.

### Genomic insights into growth-promoting mechanisms of *Penicillium melinii*

To further characterize the mechanisms underlying *P. melinii* ‘isolate 2’ ability to enhance root growth in plants, the genome was sequenced, assembled and annotated. Sequencing of genomic DNA from **‘**isolate 2’ generated 29,254,691 paired-end short reads (150 bp), resulting in an estimated depth of 301.22×. The genome assembly of **‘**isolate 2**’** consisted of 305 contigs (≥1,000 bp) with a total length of 29,091,016 bp (Figure 3A). Gene prediction identified 10,514 coding sequences (CDS), of which 87.4% (n = 9,194) were successfully annotated into functional categories (Figure 3C).

To identify potential molecular determinants underlying the PGP traits of ‘isolate 2’ such as putative cross-kingdom interactors with plant roots, the functional annotation was focused on predicting the set of proteins likely to be secreted by this strain (Figure 3B). Approximately 8% of the predicted proteome (n = 843) contained a signal peptide, while 12.95% (n = 1,362) exhibited at least one transmembrane domain. All predicted proteins with a signal peptide and without transmembrane domain were classified as secreted proteins candidates and represented 6.6% (n=696) of the whole predicted **‘**isolate 2**’** proteome. CAZymes constitute a key group of proteins involved in PGP colonization, facilitating the establishment of a close endophytic interaction with the host (Mesny et al., 2021). In the genome of ‘isolate 2’, CAZymes were identified along with other protein categories associated with plant cell wall degradation and host contact, such as proteases and putative effectors (Figure 3C). The CAZyme profile of ‘isolate 2’ was dominated by Glycoside Hydrolases (GH) and Auxiliary Activities (AA) families (Figure 3D). Among proteins with proteolytic activity, serine proteases represented the most abundant group in the genome of ‘isolate 2’ (Figure 3B). Using the EffectorP tool, putative effectors were predicted among the secreted protein candidates. In ‘isolate 2’, effectors represented 26.15% (n = 182) of the secretome (Figure 3C). This proportion is comparable to that reported for endophytic fungi and substantially lower than values typically observed in pathogenic species. EffectorP predictions indicated that 76.37% (n = 139) of these effectors are likely to function in the host apoplast, 19.78% (n = 36) in the cytoplasm, and 3.85% (n = 7), in both compartments.

To elucidate the evolutionary position of ‘isolate 2’ within the *Penicillium* genus and explore its genome-wide relationships, a phylogenomic analysis was conducted using an exceptionally comprehensive dataset of 607 *Penicillium* genomes and 100 single-copy BUSCO genes. This large-scale approach provided robust resolution of major subclades, including *Penicillium* (subclade A) and *Aspergilloides* (subclade B) (Figure 4). Although the genome of *P. melinii* has not been previously sequenced or publicly available, the phylogenomic inference for ‘isolate 2’ was fully consistent with our multi-locus phylogenetic analysis and with previously reported phylogenetic and phylogenomic frameworks for the genus (Houbraken & Samson, 2011; Steenwyk et al., 2019; Houbraken et al., 2020). Genome-based taxonomic inference placed ‘isolate 2’ within the *Exilicaulis* section, identifying *P. maclennaniae* (GCA_028827695.1), *P. atrosanguineum* (GCA_028827265.1, GCA_028827005.1, GCA_028826765.1, GCA_028826785.1), and *P. corylophilum* (GCA_018410175.1, GCA_018410145.1) as its closest relatives, with a bootstrap support of 100% (Figure 4b). This analysis enriches the genomic landscape of *Penicillium* by integrating a previously unsequenced genome and assessing its evolutionary relationships within the context of all currently published species.

**Figure 4.**
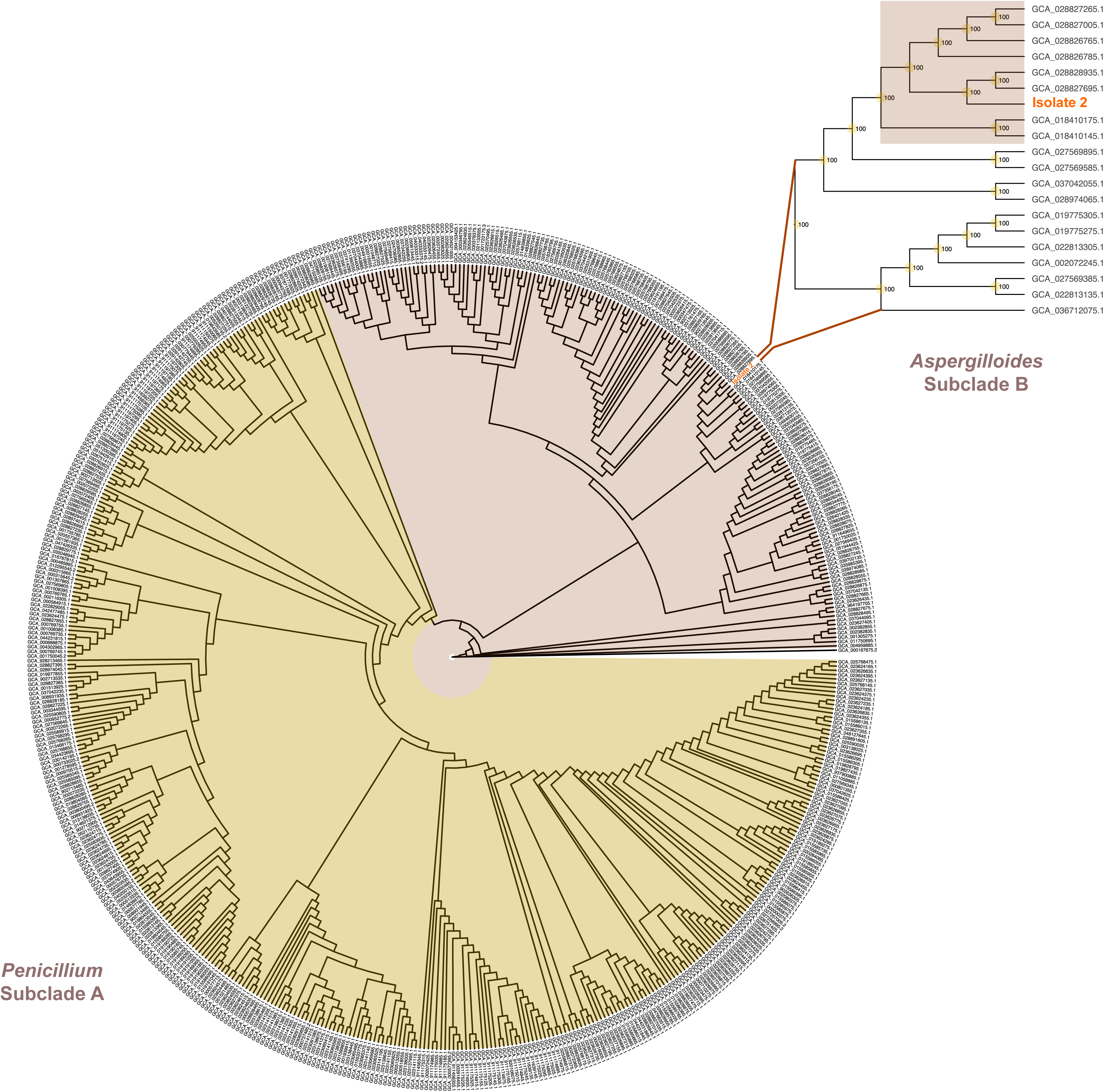
Phylogenomic placement and evolutionary relationships of ‘isolate 2’. Circular phylogenetic tree of Isolate 2 and 607 additional *Penicillium* species, inferred with IQ-TREE using 100 randomly selected BUSCO genes. The tree is rooted with *Trichoderma reesei* QM6a (GCA_000167675.2) as the outgroup. Two major subclades are highlighted: subclade A corresponds to *Penicillium*, and subclade B to *Aspergilloides*. The branch representing ‘isolate 2’ is emphasized in orange. The upper-right inset shows an enlarged view of the ‘isolate 2’ cluster within subclade B, section *Exilicaulis*. Bootstrap support values (100%) are indicated at each node, reflecting high confidence in the inferred relationships.

**Figure 5.**
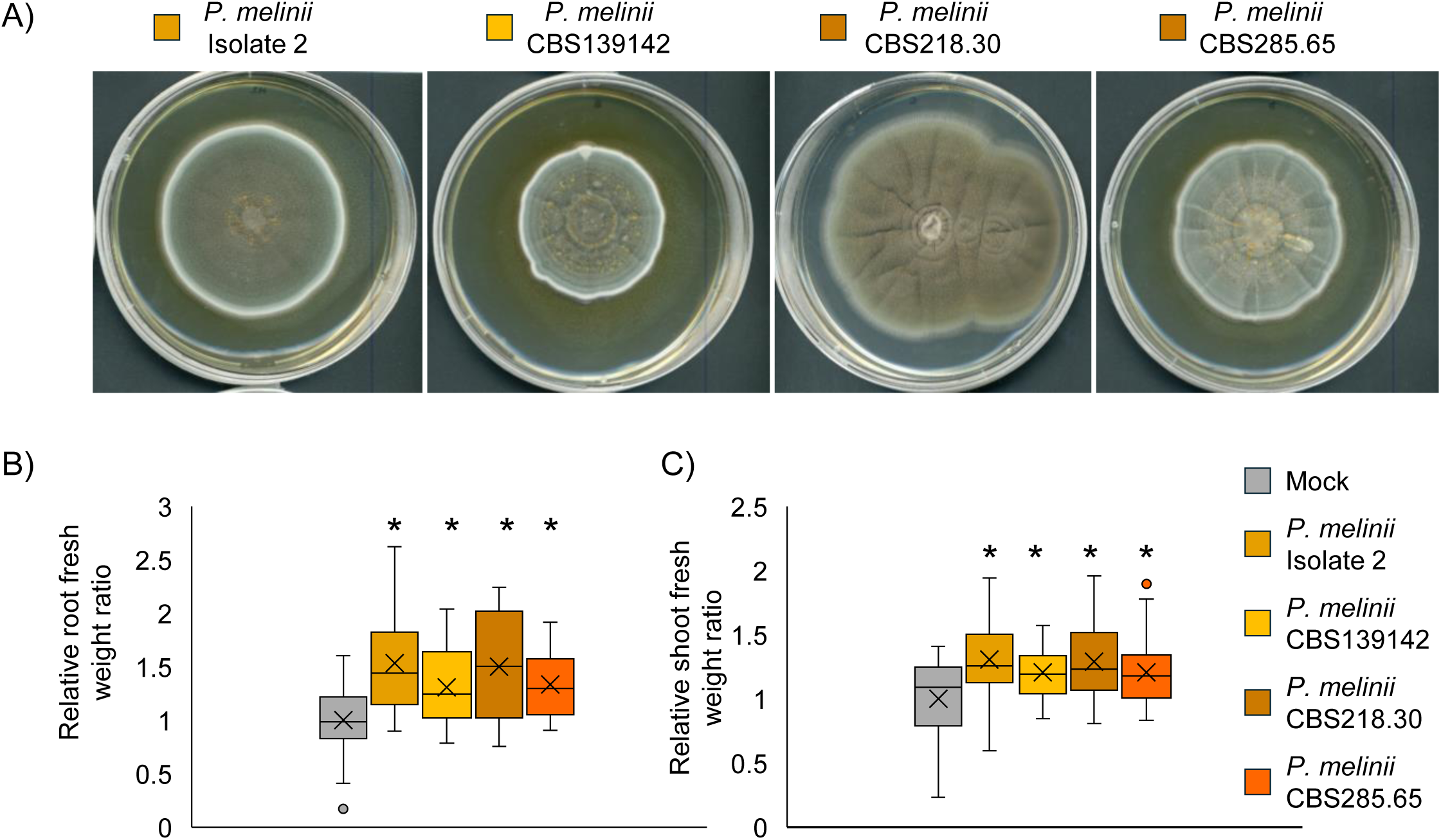
Growth-promoting effects of different *Penicillium melinii* strains. (A) Representative images of *P. melinii* strains obtained from international public repositories: CBS139142 (South Africa), CBS218.30 (USA), and CBS285.65 (England). Cultures were grown on potato dextrose agar for 10 days. (B, C) Quantification of shoot (B) and root (C) fresh weight of tomato plants 10 days post-inoculation with the respective *P. melinii* strains. Asterisks denote statistically significant differences (Student’s t-test, n > 16, P < 0.05).

### *P. melinii* induces a subtle transcriptomic response in Arabidopsis roots

To investigate the molecular mechanisms underlying *P. melinii*-enhanced LR formation, RNA sequencing on *A. thaliana* roots inoculated with the strain ‘isolate 2’ at two time points, three- and six- days post co-cultivation, was performed (Figure 6). At day 3, only 142 genes were differentially expressed (DEGs; Figure 6A). Out of these, 133 genes (93.6%) were upregulated, indicating a mild but directed transcriptional response with almost no gene repression observed. Notably, 28 of the genes induced at day 3 (21%) remained upregulated at day 6, when a total of 152 genes showed increased expression (Figure 6A–B). By day 6, however, a marked rise in gene repression was evident, with 131 genes (41.2% of DEGs) downregulated.

**Figure 6.**
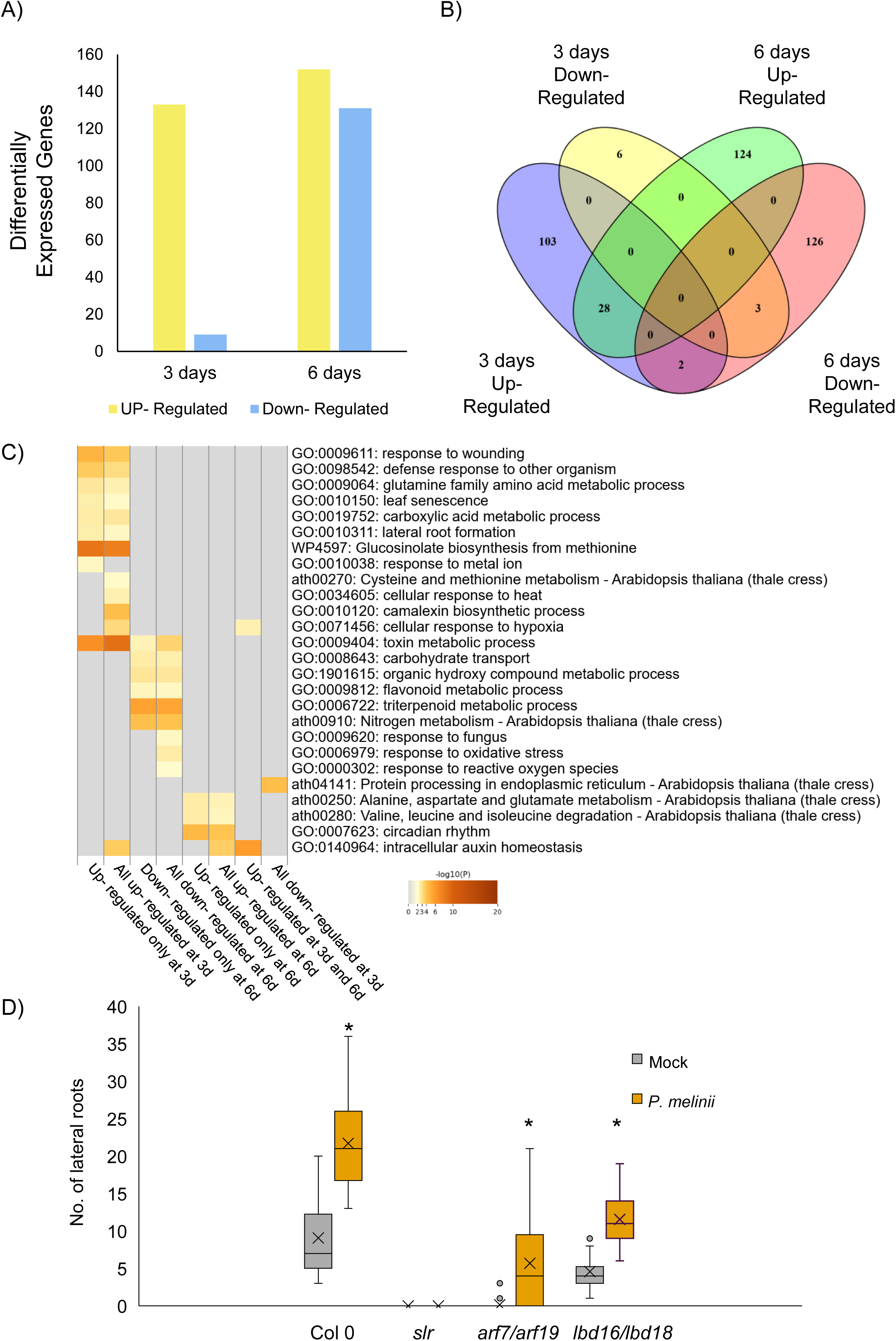
*Penicillium melinii* induces a subtle transcriptomic response in Arabidopsis roots. (A) Differentially expressed genes in Arabidopsis roots after 3 and 6 days of co-incubation with the fungus *P. melinii*. Yellow indicates up-regulation and blue down-regulation. FDR<0.05 and Fold Change ≥ 0.5 or ≤-0.5. (B) Venn diagram of the genes up- and down-regulated at 3 and 6 days. (C) Gene ontology and kegg term enrichment analysis of the groups of genes differentially expressed in the transcriptomic analysis. Only those categories overrepressented at least in one of the groups are shown. (D) Number of lateral roots in Arabidopsis wild type Col 0 and different mutants for lateral root formation inoculated with the fungus *P. melinii* after 7 days of co-incubation. Asterisks denote statistically significant differences (Student’s t-test, n > 30, P < 0.05).

Gene Ontology (GO) term enrichment analysis of these groups of genes revealed a significant overrepresentation of terms related to LR formation, including key regulators such as *LBD18* and *LBD29*, among the up-regulated genes, reinforcing *P. melinii*’s role in promoting root system development (Figure 6C). Consistent with these findings, interaction assays were conducted to assess the growth-promoting capacity of *P. melinii* in mutants impaired in key genes involved in LR formation, including *lbd16/lbd18*, *arf7/arf19* and *slr*. *P. melinii* lost its growth-promoting ability in the *slr* mutant, suggesting that this pathway is essential for LR induction during its interaction with Arabidopsis (Figure 6D). Unexpectedly, in the *arf7/arf19* double mutant, plants inoculated with the fungus still exhibited increased LR formation (Figure 6D), implying a possible redundancy among other ARFs in activating downstream targets. A similar outcome was observed in the *lbd16/lbd18* double mutant, where inoculation with *P. melinii* resulted in a clear enhancement of LR formation similar to that found in Col-0 (Figure 6D).

As expected, GO categories related to plant interaction with another organism, such as ‘Response to other organism’, ‘Response to wounding’, ‘Glucosinolate biosynthesis’, and ‘Camalexin biosynthetic process’ were enriched among upregulated genes at day 3. Interestingly, this response to the interaction with the fungus appeared to be attenuated by day 6, with terms like ‘Response to fungus’ and ‘Response to oxidative stress’ enriched among the downregulated genes (Figure 6C). Interestingly, genes involved in ‘Triterpenoid metabolism’ were repressed at day 6, with strong downregulation observed in the thalianol and marneral biosynthetic clusters, which showed no significant changes at day 3 (Figure S2). Finally, genes associated with ‘Auxin homeostasis’ were consistently upregulated at both time points, suggesting a sustained modulation of auxin-related pathways during the interaction.

### *P. melinii* induces a subtle hormonal response in Arabidopsis roots

In addition to transcriptionally altering the root response, endophytes are known to modify the hormonal balance of the plant (Egamberdieva et al., 2017). To assess whether *P. melinii* influences hormonal signalling in *A. thaliana* roots, hormone-related metabolites were profiled using the method described by Simura et al. (2018) for samples collected from mock- and *P. melinii*- inoculated plants at 3-, 6-, and 12- days post-interaction (Figure 7A). Consistent with the modest transcriptomic changes observed, alterations in hormone levels were limited. At day 3, only phaseic acid (PA), a degradation product of abscisic acid (ABA), was significantly elevated in *P. melinii*-treated roots (Figure 7A). By day 6, most differentially accumulated metabolites were found in reduced quantities. Notably, intermediates involved in indole acetic acid and glucosinolate biosynthesis, such as tryptophan (TRP), tryptamine (TRA), indole-3-acetonitrile (IAN), and indol-3-acetamid (IAM), were less abundant in inoculated roots, aligning with the transcriptomic data showing upregulation of camalexin and glucosinolate-related genes at day 3 but not at day 6. In addition to auxin-related compounds, several cytokinins-including isopentenyladenine riboside (iPR), isopentenyladenine-7-glucoside (iP7G), cis-zeatin riboside-O-glucoside (cZROG) and cis-zeatin-O-glucoside (cZOG), were also under-accumulated in *P. melinii*-treated roots. Furthermore, key stress-related hormones such as salicylic acid and jasmonate-isoleucine conjugates were found in lower concentrations at days 6 and 12, suggesting a suppression of the plant response to these biotic interactions. This observation is consistent with transcriptomic data indicating an initial activation of stress-related genes on day 3, followed by downregulation at day 6.

**Figure 7.**
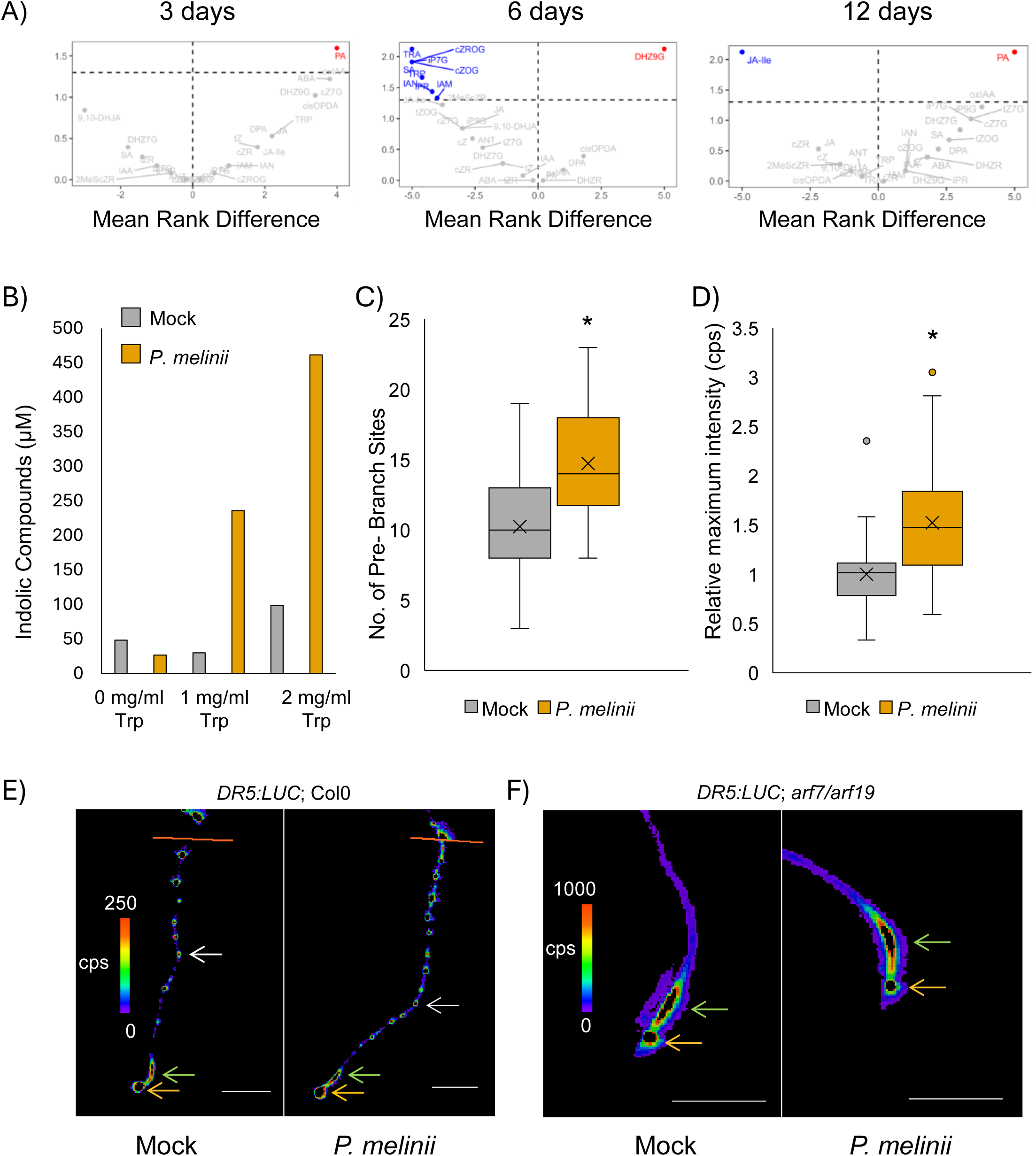
*Penicillium melinii* induces a subtle hormonal response in Arabidopsis roots. (A) Differentially accumulated precursor compounds of the different hormones in Arabidopsis roots inoculated with *P. melinii* after 2, 6 or 12 days of interaction. Each point in the volcano plot represents a plant hormone metabolite, positioned according to the rank difference between Mock and *P. melinii* treatments on the x-axis and the corresponding p-value from the Wilcoxon-Mann-Whitney U test on the y-axis. Negative x-axis values indicate downregulation, while positive values indicate upregulation. P-values are shown as, log_10_-transformed values; metabolites above the horizontal dashed line exhibit statistically significant differences at the 5% significance level. (B) Quantity of indolic compounds produced by *P. melinii* when incubated with Salkowski reagent and increased quantities of tryptophan. (C) Number of pre-branch sites induced by *P. melinii* in *DR5:LUC* plants after 3 days of interaction. Asterisks denote statistically significant differences (Student’s t-test, n > 58, P < 0.05). (D) Relative intensity of the luminescence in the OZ of *DR5:LUC;arf7/arf19* plants treated or not with *P. melinii* Asterisks denote statistically significant differences (Student’s t-test, n > 37, P < 0.05). (E-F) Representative pictures of the auxin signalling indicated by *DR5:LUC* promoter in Col-0 (E) and *arf7/arf19* (F). White arrows indicate pre-branch sites. Green arrows indicate the OZ. Orange arrows indicate the root apical meristem.

Despite the hormonome-related results obtained from plants inoculated with the fungus, the Salkowski reagent assay demonstrated that *P. melinii* ‘isolate 2’ cultured in liquid medium produces indole-derived compounds and releases them into the surrounding medium (Figure 7B). Auxin signalling was subsequently analysed using the *DR5:LUC* reporter line, which allows the analysis of the auxin response and the quantification of PBS, the precursors of LRs, originating from the OZ (Figure 7C–E). Consistent with the growth-promoting capacity of *P. melinii* and the hormonome-related data, roots exposed to the fungus displayed a significantly higher number of PBS compared with mock plants, although no substantial increase in ectopic auxin signalling was observed. Taken together, these findings suggest that *P. melinii* specifically promotes LR formation without globally activating auxin signalling (Figure 7C–E). In the *arf7/arf19* mutant line, auxin signalling in the root does not oscillate within the OZ; therefore, measuring luminescence intensity in this region provides an estimation of auxin signalling strength (Figure 7D–F). Notably, plants treated with *P. melinii* exhibited increased auxin signalling intensity in the OZ compared with mock plants (Figure 7D–F), suggesting that, although signalling is not ectopically induced, it may be locally enhanced in regions where it drives LR formation during the interaction *with P. melinii*.

## Discussion

Understanding how beneficial endophytic fungi influence root system architecture is critical for improving plant resilience and productivity in sustainable agriculture. In this work, it is provided a comprehensive characterisation of the interaction between *P. melinii* and plants, revealing how this fungus promotes LR formation, resulting in an enlargement and remodelling of the root system. Our findings integrate phenotypic, transcriptomic, hormonal, and genetic evidence to show that *P. melinii* employs a finely tuned strategy that balances growth and the use of secondary metabolites, positioning this species as a promising candidate for biostimulant applications in crops.

The strain ‘isolate 2’ from *P. melinii* was selected for its effect on root growth from an endophytic fungi collection from Arabidopsis (García et al., 2013). Specifically, ‘isolate 2’ was isolated from the roots of an Arabidopsis wild specimen growing in the countryside of Polan (Toledo, Spain; García et al., 2013). Previous studies investigating fungal diversity in plant-associated soils have reported *P. melinii* as one of the most abundant species in boreal mixed-wood forest soils in Quebec (De Bellis et al., 2007) and in the rhizosphere of the halophytic grass *Aeluropus littoralis* (Tarroum et al., 2021). In the latter study, mycelial growth or cell-free filtrates of *P. melinii* were shown to enhance root length and biomass of tobacco seedlings grown on plates or in nutrient solution for 3–4 weeks (Tarroum et al., 2021). However, the detailed characterisation of this interaction and its influence on root system architecture has remained largely unexplored. Here, it is provided a comprehensive analysis of how this fungus modulates root development in Arabidopsis, notably increasing the number of LRs (Figure 1) and altering the root clock mechanism (Moreno-Risueno et al., 2010), as demonstrated by the measurements of a stronger auxin signalling response within the OZ and a higher number of PBS (Figure 7).

Not only the beneficial effect of *P. melinii in vitro* using the model plant Arabidopsis is demonstrated but also extended these findings to crop species such as tomato and quinoa grown under greenhouse conditions, and, importantly, translated this knowledge to plants cultivated in agricultural fields (Figure 2). Furthermore, it is demonstrated that this positive effect is not restricted to the ‘isolate 2’ described by García et al. (2013) but can be generalised to the species as a whole, as three additional strains from diverse geographical origins exhibited the same effect on tomato plants (Figure 5C-D). The persistence of the growth-promoting effect of *P. melinii* both *in vitro* with model plants and in soil with crops suggest a remarkable robustness of this interaction, a feature also observed for other beneficial fungi that consistently enhance plant growth across controlled and field conditions (Adedayo and Babalola, 2023). The use of rhizotrons to grow tomato and quinoa enabled us to better characterise root growth over time, in contrast to the experiments conducted in pots or liquid culture (Tarroum et al., 2021). The progression from controlled experiments to real agricultural settings, underscores the potential of *P. melinii* as a sustainable strategy to enhance root development and improve crop performance. It also highlights this study as a clear example of the transferable capacity of findings obtained *in vitro* (with subtle changes at morphological, transcriptomic and hormonal levels) to practical field applications (where there is an impact on plant fresh weight and productivity), as it has also been shown with other fungal endophytes of Arabidopsis (Hiruma et al., 2016; Díaz-González et al., 2020.

The use of a GFP-expressing transgenic strain demonstrated that, under our experimental conditions, *P. melinii* established a profuse colonisation of the root surface, particularly at the epidermis and cortex (Figure 1), while no hyphae were detected within the vascular cylinder. This observation, together with the previously reported growth-promoting effect of cell-free filtrates of *P. melinii* (Tarroum et al., 2021), strongly suggests that the fungus exerts its influence through secreted metabolites, likely in these two cell layers, although additional work will be needed. Similar surface colonisation and growth-promoting properties of fungal exudates have been described for a GFP-expressing strain of the closely related endophyte *P. citrinum* in *Brassica rapa* var. *parachinensis*, where growth promotion was most likely mediated by fungus-derived gibberellins (Gu et al., 2023). Indeed, the sequencing and analysis of *P. melinii* ‘isolate 2’ genome reveal genes related to the biosynthesis of secondary metabolites and of putative secreted proteins (Figure 3). The relatively low proportion of predicted effectors within the *P. melinii* secretome (26%) is consistent with values reported for endophytic fungi and contrasts sharply with pathogenic species, where effector expansion is a hallmark of virulence (Hacquard et al., 2016; Jia et al., 2023; Moonjely and Trail, 2025). Pathogens often deploy large repertoires of lineage-specific effectors to suppress host immunity and manipulate plant physiology aggressively, with effector-like proteins constituting up to half of the secretome in some biotrophic fungi (Chakraborty et al., 2023). This pattern suggests that *P. melinii* relies on a strategy of metabolic exchange and signalling rather than pathogenic interference, reinforcing its role as a beneficial symbiont and supporting its potential application as biostimulant. According to EffectorP predictions, 76.37% of the effectors are likely to act in the host apoplast, 19.78% in the cytoplasm, and 3.85% in both compartments. This distribution is consistent with previous observations that filamentous fungi predominantly reside in the host apoplast during the early stages of colonisation (Rocafort et al., 2020).

This study reports the first genome of *P. melinii* ‘Isolate 2’, offering a detailed gene catalogue and functional annotation. Our analyses reveal a secretome dominated by apoplastic effectors, consistent with a symbiotic lifestyle, and identify candidate genes linked to plant growth promotion. The present work establishes an unprecedented and rigorous taxonomic framework integrating both loci-based and genome-based approaches, the latter encompassing all currently available *Penicillium* genome assemblies. Considering the remarkable diversity of *Penicillium* genus and the persistent challenges in resolving its taxonomy (Mahmoud et al.,2016; Visagie at al., 2016), the taxonomic classification of *P. melinii* presented here represents a valuable resource for advancing systematic knowledge and functional insights into this ecologically and industrially important group of fungi.

To understand the basis of plant-growth promotion facilitated by *P. melinii*, transcriptomic and hormonal responses of Arabidopsis roots inoculated with the fungus, were evaluated. This analysis reveals a transcriptomic and hormonal pattern typical of beneficial endophytic associations, where the plant balances plant growth and plant response to other organisms to enable compatibility. Our data show that, three days post-inoculation, the response is subtle yet directional, with 93.6% of differentially expressed genes (DEGs) upregulated and Gene Ontology categories linked to ‘response to other organism’, ‘glucosinolate biosynthesis’, and ‘camalexin biosynthesis’. This initial response aligns with previous studies on plant-endophyte interactions, which report a transient activation of plant response to microorganism pathways followed by attenuation to consolidate symbiosis (Hardoim et al., 2015). Likewise, early induction of indolic pathways (camalexin and glucosinolates) followed by their decline reflects a transition from resistance to compatibility, which may depend on the nutritional status of the plant. Hacquard et al. showed that responses related with indole-glucosinolates and ethanol metabolic processes are activated in Arabidopsis roots colonized by *Colletotrichum tofieldiae* under sufficient phosphate levels, but not under phosphate starvation (Hacquard et al., 2016). Moreover, intriguingly, metabolites such as camalexin and thalianol, deregulated by *P. melinii* in our transcriptome, have recently been implicated in the regulation of root system architecture and specific microbiome recruitment (Bai et al., 2021; Huang et al., 2019; Serrano-Ron et al., 2021). Alterations in specialised metabolites may facilitate endophyte colonisation by shaping plant–microbe interactions (Huang et al., 2019). These compounds, traditionally associated with a function in the response to biotic interactions, can act as selective filters, enabling certain endophytes to tolerate them. This adaptive capacity likely promotes mutualistic relationships and influences the assembly of root-associated microbial communities (Huang et al., 2019).

In parallel, the transcriptomic analysis showed a sustained up-regulation of auxin homeostasis-related genes, but no ectopic activation through the whole root system was observed using the reporter line *DR5:LUC*. Using this reporter line revealed a significant increase in the number of PBS and higher luminescence intensity in the OZ. Similar localized auxin responses have been described in the interaction between the endophytic fungus *Serendipita indica* and Arabidopsis, where alterations in auxin transport and conjugation generate local auxin maxima that promote root growth (González Ortega-Villaizán et al., 2024). The production of indolic compounds by *P. melinii* was demonstrated, in agreement with previous reports describing the ability of many endophytes, including *Trichoderma* spp. and *Falciphora oryzae*, to produce auxin-like molecules (Contreras-Cornejo et al., 2009; Sun et al., 2020). Beyond auxins, numerous endophytic genera are also capable of synthesizing cytokinins and gibberellins. For example, within *Trichoderma*, several species produce auxins that reshape the root system (Contreras-Cornejo et al., 2009), up to 11 forms of cytokinins detected across 22 strains (Bean et al., 2022), and gibberellin A3 isolated from culture filtrates (Kamalov et al., 2018). It is important to note that although the Salkowski assay confirmed the presence of indolic compounds in *P. melinii* cultures, this method is not specific for IAA, and validation by LC–MS/MS is required for precise identification. Indeed, in our hormone profiling, IAA did not accumulate to higher levels in inoculated roots, suggesting that the observed root phenotypes may result either from localized auxin modulation or from additional hormone-related mechanisms triggered by the endophyte.

An unexpected finding was the persistence of LR promotion in the *arf7/arf19* double mutant, key genes in the ARF–LBD module regulating LR development. The persistence of this effect in double mutants points to functional redundancies or alternative routes converging on downstream effectors such as *LBD29* or *PRH1*, both implicated in LR formation. Notably, *LBD29* was significantly upregulated in roots treated with *P. melinii*, while *PRH1* displayed a clear tendency towards upregulation, although not reaching statistical significance (Okushima et al., 2007; Zhang et al., 2020). Furthermore, *LBD18,* whose overexpression can rescue LR formation even in the absence of *ARF7* and *ARF19,* was up-regulated during the interaction with *P. melinii* (Lee et al., 2019). This observation is consistent with recent evidence that plant-associated microbiota induce LR formation in the *arf7/arf19* mutant, activating root branching via mechanisms independent of classical auxin signalling, including ethylene-mediated routes (Gonin et al., 2023). It has also been reported that the colonization by *Sinomonas gamaensis* can remodel the *IAA14-ARF7/ARF19* interaction independently of *TIR1/AFB2* (Fu et al., 2025). Similarly, LR promotion in the *lbd16/lbd18* mutant suggests that *P. melinii* may activate other LBD-family members or employ additional signals to trigger the LR formation programme.

Taken together, our findings indicate that *P. melinii* promotes LR formation through a multifactorial strategy that begins with an initial activation of plant responses to microorganism followed by their attenuation to enable compatibility, while simultaneously fine-tuning auxin signalling locally within the OZ rather than inducing global hyperactivation. This targeted modulation appears to work in concert with a potential bypass of the canonical ARF7/ARF19 pathway through downstream redundancies, ensuring that LR initiation can proceed even when key regulators are impaired. In addition, the interaction involves discrete metabolic and hormonal reprogramming, including adjustments in specialised metabolite pathways and phytohormone profiles, which collectively create a physiological environment conducive to root system remodelling and sustained symbiosis.

## Acknowledgments

This work has been funded by MCIN/AEI/ 10.13039/501100011033 and FSE through the projects: RYC2021-030913-I and PID2022-141938OA-I00 to JC; PID2020-113479RB-I00 to JCP; and ‘Severo Ochoa Program for Centres of Excellence in R&D’ from the ‘Agencia Estatal de Investigación’ of Spain CEX2020-000999-S grant to the CBGP. ALE is supported by a grant ‘PRE2021-098261’. This work was also supported by the Johannes Amos Comenius Programme via the project “TowArds Next GENeration Crops (TANGENC)” (No. CZ.02.01.01/00/22_008/0004581) from the Ministry of Education, Youth and Sports of the Czech Republic. *LTi6b-mCherry* line and pLM2 plasmid were kindly provided by Dr. Alexis Maizel’s laboratory (Germany) and Dr. Andrea Sánchez-Vallet laboratory (Spain), respectively.

## Author contributions

CC, LU, JN, AVM, CCBL, EAO, MAMR, MR, SS, JCP and JC planned and designed the research. LGM, IDA, CC, AL PFC, IP, AP, MAMR, MR and JC performed the experiments. LLG, IDA, GMG, SGS, FB, LU, AVM, EAO, SGB, JHC, SS, JCP and JC analysed the data. LGM, IDA, CC, SGS, FB, PFC, MR, JCP and JC wrote the manuscript. CC, SGB, JHC, SS, JCP and JC supervised the research. LU, JN, MAMR, JHC, SS, JCP and JC acquired the funding. All authors revised the final version of the manuscript.

