## Supplementary material for "Root growth promotion by *Penicillium melinii*: mechanistic insights and agricultural applications": Figure S1

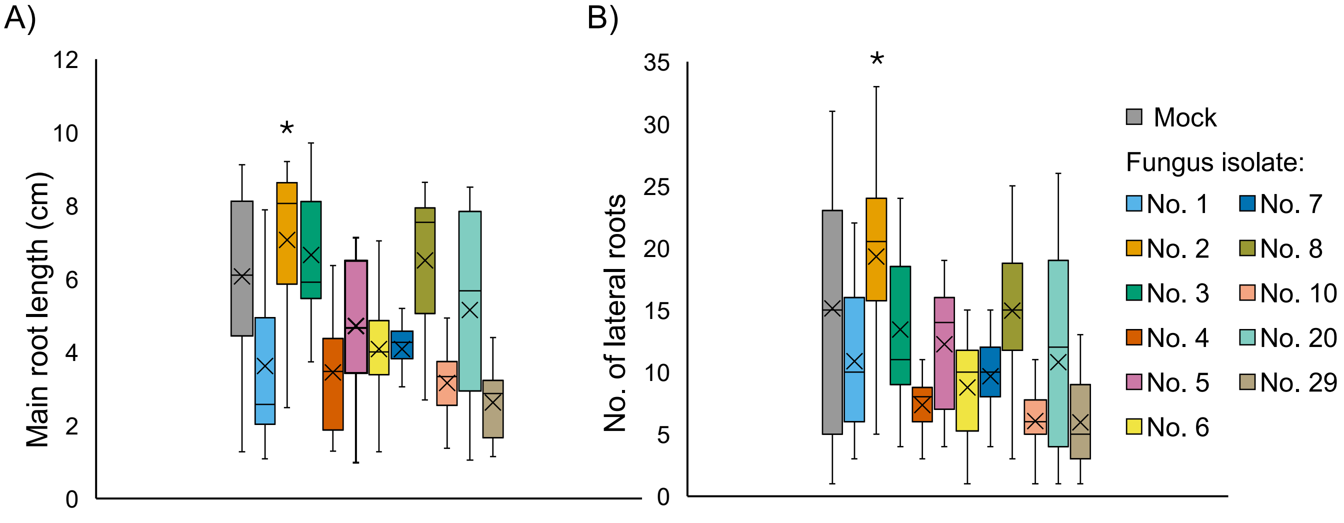


**Figure S1. Screening of Arabidopsis fungal endophytes for root growth promotion.** (A) Primary root length and (B) number of lateral roots measured seven days after incubation of Arabidopsis seedlings with different fungal isolates. Measurements were taken from the root section that developed on the fungal-inoculated plate after seedling transfer. Asterisks indicate statistical significance (Student's t-test, n > 13, P < 0.05).
