## Supplementary material for "Root growth promotion by *Penicillium melinii*: mechanistic insights and agricultural applications": Figure S2

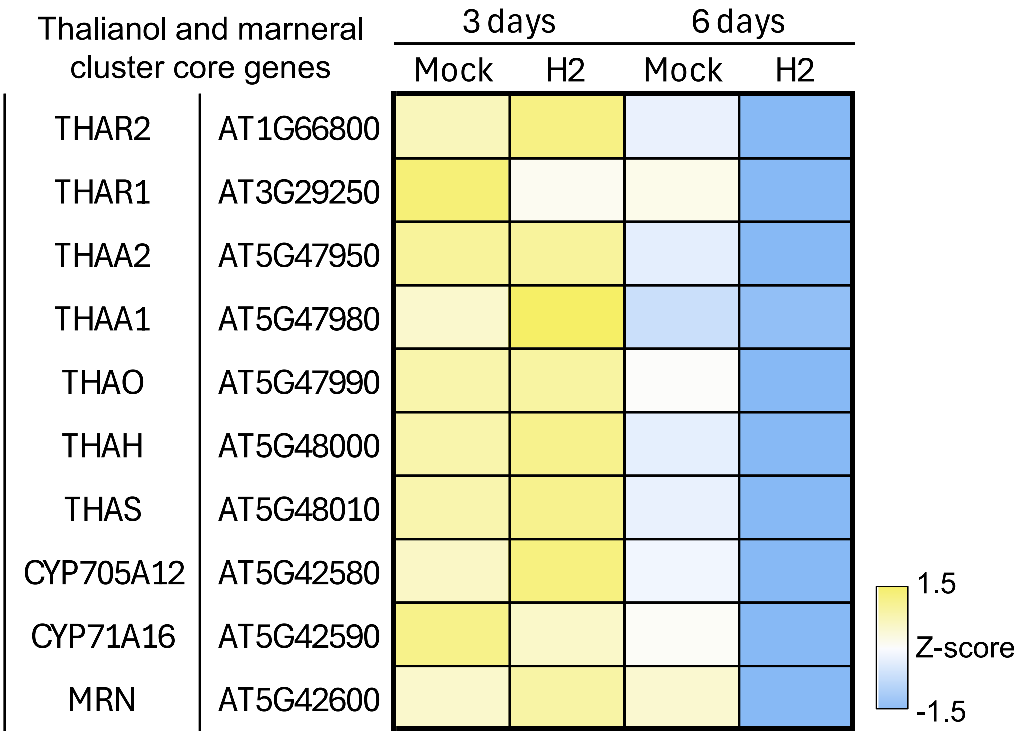


**Figure S2.** Heatmap of Z- score values of the expression of those genes participating in the thalianol and marneral synthesis.
